# Methane production from hydrocarbons by consortia dominated by ANME archaea

**DOI:** 10.1101/2024.10.30.621140

**Authors:** Natalie Sarno, Andrew Montgomery, Guang-Chao Zhuang, Brennan R. Martinez, Valerie De Anda, Samantha B. Joye, Brett J. Baker

## Abstract

Understanding microbial production and consumption of methane, a potent greenhouse gas in the atmosphere, is critical for bridging knowledge gaps in global carbon cycling. In anoxic environments, methane is known to be produced through hydrogenotrophic, acetoclastic or methylotrophic mechanisms. Methane production from hydrocarbons may also be important, especially in hydrocarbon-rich environments, like the Gulf of California, but the mechanism of this hydrocarbonoclastic methanogenesis remains unclear. The activity of consortia of anaerobic methane oxidizing (ANME) archaea and bacteria limits the release of methane to the atmosphere by consuming methane in anoxic environments globally. Here we used isotopic-labeling to track the conversion of hydrocarbons (hexadecane and naphthalene) to methane in enrichments from hydrothermally impacted, hydrocarbon-rich sediments from the Gulf of California. Methane was produced directly from hexadecane and naphthalene, in both the presence and absence of sulfate. We reconstructed metagenomic assembled-genomes (MAGs) from these experiments which revealed a mixture of bacteria dominated by Desulfobacteriota and Bacteroidota, and archaea dominated by Aeinigmarchaeota, Thermoplasmatota, and ANME group 2c. The ANME-2c were the only MAGs that encoded methyl coenzyme M reductases (McrA) and complete Wood-Ljungdahl pathways (WLP). This suggests that ANME-2c archaea may be involved in the production of methane along the seafloor, and that our understanding of the roles of these globally important microbes is not yet fully appreciated.

---

Methane, a potent atmospheric greenhouse gas, is generated predominantly in anoxic environments through the breakdown of organic matter. The majority of this methane is then consumed by methanotrophs, including anaerobic methane oxidizing (ANME) archaea^1–4^. Recently, it was suggested that ANME archaea are capable of reversing the methanotrophic pathway to produce methane^5,6^. There are also indications of methane production from hydrogen and carbon dioxide (CO_2_) in enrichments containing ANME-1^7^. Yet the direct involvement of ANME in methanogenesis remains to be confirmed. Another novel methane producing metabolic pathway in anoxic environments is the degradation of hydrocarbons via a divergent “mcr-like” homolog called alkyl-coenzyme M reductase (Acr)^8^. This particular Acr activates complex hydrocarbons (*e*.*g*., hexadecane or hexadecylbenzene) into an alcohol via a coenzyme M intermediate. Ultimately, Acetyl-CoA is routed to the Wood-Ljungdahl pathway (WLP) and complete methanogenesis pathways^8^. Hydrocarbons, organic molecules composed of carbon and hydrogen atoms, are generated by thermocatalytic cracking of organic matter and are abundant in hydrothermally-heated and cold seep sediments in the Gulf of California. Oil is produced in subsurface environments elsewhere but the petroleum formation window is much shallower and the characteristics of hydrothermal oil are distinct^9^. Typically, community degradation of hydrocarbons leads to production of carbon dioxide, however, one organism (an Euryarchaeota) has been demonstrated to convert hydrocarbons directly to methane via Acr activation^8^. Here we show that enrichments dominated by ANME archaea are capable of methanogenesis.

## Results

To characterize microbial mediated hydrocarbon and methane cycling in the deep-ocean we examined hydrothermally altered sediments from the Cathedral Hill, a hydrothermally-impacted site 2,000 m below the sea surface in Guaymas Basin (Gulf of California). Methane and dissolved organic carbon (DOC) concentrations increased with depth to a maximum of 885 µM and 3.7 mM respectively and anaerobic oxidation of methane (AOM) rates followed a similar trend (Figure 1). Porewater sulfate concentrations decreased with depth and coincided with sulfide accumulation and sulfate reduction rate up to 4 mM and 7.7 nmol cc^-1^ d^-1^ respectively (Figure 1). Below 7.5 cm, AOM rates were substantially higher than sulfate reduction rates indicating a decoupling of AOM and sulfate reduction. The sediment surface was covered with an orange *Beggiatoa* mat and the sediment temperature ranged from 4°C at the sediment water interface to 55°C at 10 cm. We amended these sediments with crude oil to a final concentration of 0.12% w/v. The accumulation of sulfide was used as a proxy for sulfate reduction. After one transfer to fresh media (with a 1:5 dilution) and persistent sulfide production, methane production rates were measured using radiotracer assays.

**Figure 1.**
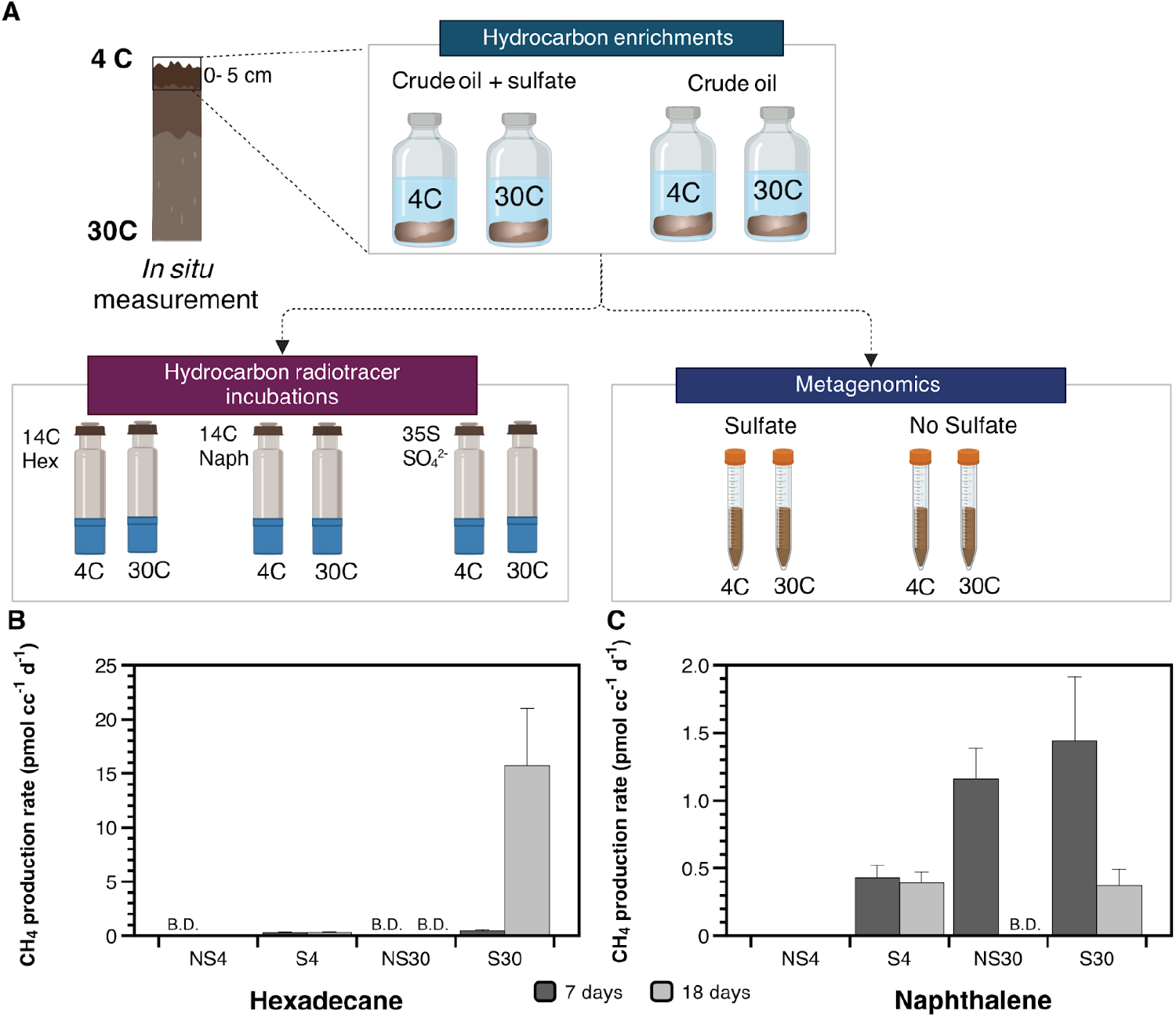
Shifts in geochemistry downcore in the Gulf of California sediments and enrichments during methane-production. (A) Workflow of mesocosm enrichments and subsampling for genomic and rate measurements. The 0-5 cm horizons from two cores were homogenized and divided equally by mass. Each portion of sediment was then allotted into artificial seawater medium, one with sulfate (28 mM final concentration) and one with no sulfate added. Four replicate serum vials were made for each treatment (+/-SO_4_^2-^; 4°C and 30°C) and one of each was an autoclaved kill control. All serum vials were amended with crude oil to a final concentration of 0.15% v/v. After sulfide production was observed, subsamples for genomic and radiotracer rates were taken. (B and C) Methane production rates from (B) hexadecane and (C) naphthalene after ∼70 days enrichment with sediment from Cathedral Hill and crude oil. Methane production rate incubations were incubated for 7 (light gray) and 18 (dark gray) days NS = no sulfate added; S = 28 mM initial sulfate concentrations; 4 = 4°C incubation temperature; 30 = 30°C incubation temperature. B.D. = below detection limit.

Surprisingly, methane was produced directly from hexadecane and naphthalene, in the presence and absence of sulfate, at rates up to 1.4 ± 0.4 and 15.7 ± 5.2 pmol cc^-1^ d^-1^ for each substrate, respectively. The highest rate was observed in the sulfate treatment at 30°C, suggesting that sulfate was not an inhibitor of this mode of methanogenesis. In fact, methane production from hexadecane could not be detected in the treatments without sulfate addition indicating sulfate may even play a role in hexadecane degradation during methanogenesis in these communities. On average, rates were greater from naphthalene than hexadecane and higher temperatures had an apparent effect on naphthalene degradation. Specifically, methane production from naphthalene was greater than 1 pmol cc^-1^ d^-1^ at 30°C and was below 0.5 pmol cc^-1^ d^-1^ or below detection at 4°C.

We obtained 64 Gb of short-read shotgun genomic data from these enrichments to understand the diversity and metabolisms organisms may be able to produce methane from hydrocarbons. These data were assembled and binned into 93 metagenome assembled genomes (MAGs) that were over 50% complete (<10% gene redundancy). In order to determine the identity of these genomes we used the Genome Taxonomy Database (GTDB) and generated phylogenomic trees using a common set of 37 marker proteins^10^ (Supplementary Figure 1). The dominant phyla belong to Caldatribacteriota, Bacteroidota, Desfulobacteriota, and Halobacterota (Supplementary Table 3). Among these Halobacterota were 2 MAGs belonging to ANME group 2c (Supplementary Figure 1). CoverM^11^ was used to calculate the relative abundance of all 93 members within the communities from both the original environmental core and all enrichment samples (n=7) (Figure 2). The ANME-2c genomes, CH_155 and CH_159, were found to be the 3rd and 4th most abundant community members from the environmental core respectively (Supplementary Table 3).

**Figure 2.**
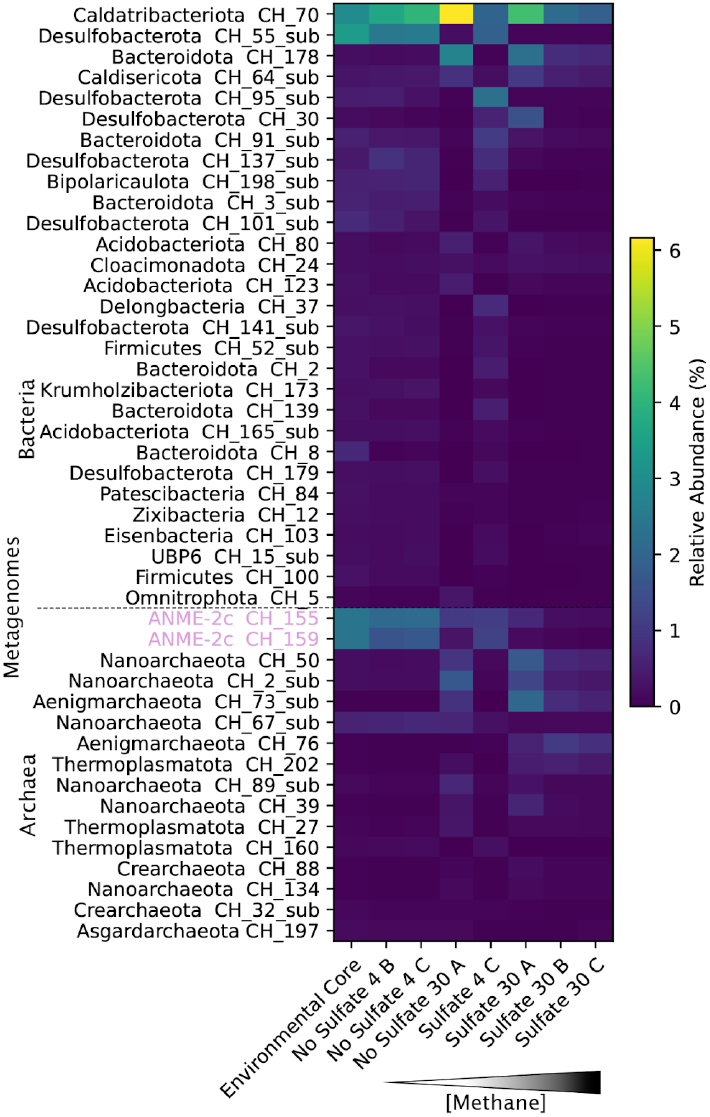
Relative abundance of the dominant enrichment microbial community members across samples. This heatmap was generated by calculating the relative abundance of the entire enrichment community (93 metagenomes) from the original environmental sediment core and across each treatment (n=7) with CoverM v0.6.1^11^. To better visualize the data, any bacterial MAG that was <0.2% relative abundance in the environment was excluded from this plot (n=48 out of 77 total bacterial MAGs). The output from CoverM was processed and visualized in Python v.3.12.0^12^ with the pandas v.2.1.1^13^, numpy v.1.26.0^14^, and matplotlib v.3.8.0^15^ libraries. Metagenomes above the dashed line are bacteria and those below are archaea. The two ANME-2c are highlighted in pink. Enrichment samples are organized from left to right by increasing methane concentrations.

We identified seven copies of the key enzyme for methane production methyl-coenzyme M reductase (McrA) in the enrichments (Figure 3A). Two of these were associated with the assembled ANME-2c MAGs, CH_155 and CH_159. The other five McrA were not binned into genomes due to lower coverage. Four of the five unbinned McrA are likely not involved in significant methane production in these enrichments because they had <6X coverage while the ANME-2c McrAs had 67X and 126X coverage respectively. The four unbinned McrAs are from Methanotrichales (6X), Methanomicrobiales (6X), Methanofastidosa (6X), and ANME group 1 (5X). The fifth unbinned McrA also belonged to ANME-2c according to BLAST^16^ with 24X coverage.

**Figure 3.**
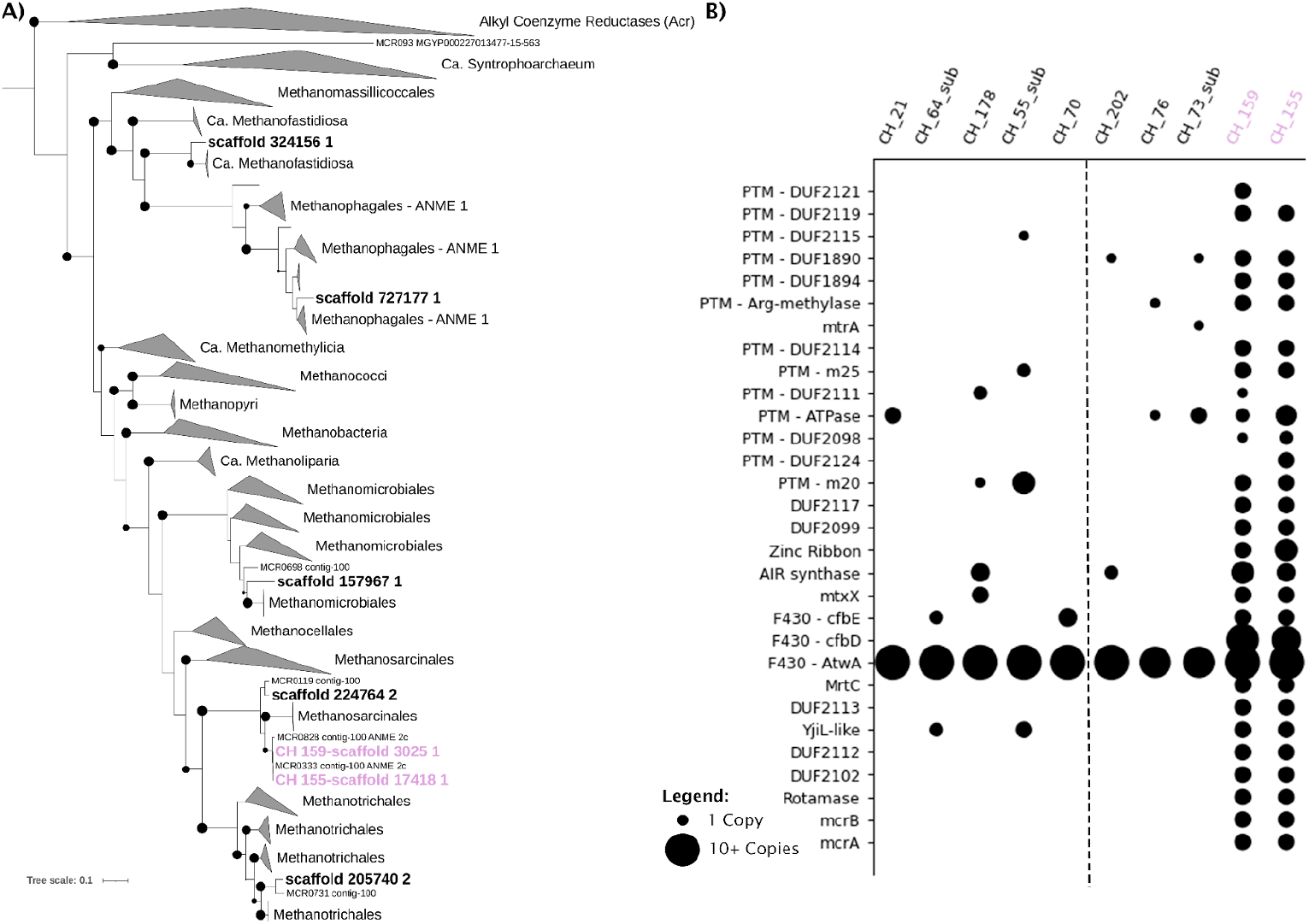
The phylogenetic position of ANME-2c McrA proteins and a bubble plot of highly conserved methanogenesis marker genes of the top 5 most abundant archaea and bacteria from the pooled assembly data. **(A)** The phylogeny on the left was generated using 1,581 McrA reference sequences^18^ for Diamond BlastP^20^ searches. There were only two McrAs found out of the 93 MAGs. Both McrAs were from ANME 2c MAGs (highlighted in pink). The assembled reads were also searched for the 1,581 McrA reference sequences as aforementioned and 5 McrAs were identified in the unbinned assembly (scaffolds in bold text). The outgroup are the alkyl coenzyme M reductase (Acr) or “mcr-like” sequences. **(B)** The bubble plot was generated with raw counts of marker gene copy numbers using Python v.3.12.0 with the pandas v.2.1.1, numpy v.1.26.0, and matplotlib v.3.8.0 libraries. Each column is an enrichment MAG while each row is a methanogenesis marker gene of interest^21^. Any markers that did not have any MAGs with copies were left out of this plot. Thus, this plot contains 30 out of the 38 markers. To the left of the dashed line are the 5 most abundant bacteria (least to most abundant left to right) followed by the 5 most abundant archaea to the right of the line (most to least abundant left to right). Smaller dots correspond to fewer copies of a gene while larger dots correspond to more copies. The two ANME-2c shown in pink (CH_155 and CH_159) were the only community members with a majority of the methanogenesis marker genes. Methanogenesis marker gene copy numbers for all 93 MAGs and all 38 markers is available in the supplementary information (Supplementary Table 5).

We searched for 38 well conserved methanogenic marker genes in these MAGs^17^. CH_155 contained 26 of these core marker genes and CH_159 contained 27 (Figure 3B). None of the other MAGs from the enrichment samples contained more than seven of these genes (Supplementary Tables 4 and 5). This suggests that ANME are the only members of these enrichments that are capable of producing methane at the levels we detected. ANME-2c MAGs encode both alpha and beta *mcr* subunits (*mcrAB*). These two MAGs also contain 12 of the 13 markers involved in post-translational modification (PTM) as well as several markers conserved in methanogens with as yet unidentified function^17^. ANME MAGs are missing three methanogenic pathway steps F_420_H_2_ oxidation via the F_420_H_2_:methanophenazine oxidoreductase complex (*fpo*) or F_420_-reducing hydrogenase (*frh*), CoM-SH/CoB-SH oxidation via heterodisulfide reductases (*hdrDE*), and the second to last step of methane production via H_4_MPT:coenzyme M methyltransferase (*mtr*). These steps have previously been shown to be missing from several ANME-2c genomes^18^. Other methanogenic archaea have also been shown to use the Rnf complex and *hdrD* instead of *fpo*^*8*^ and the *mtr* operon was removed from a methane producing Methanosarcina, yet the organism retained a functional methanogenic pathway^19^. Therefore, the three missing steps of the ANME-2c methanogenic pathway may not be essential for methane production.

Nine of the Desulfobacterota MAGs contain dissimilatory sulfite reductases (DsrAB) indicating they were involved in the sulfate reduction observed in the enrichments (Supplementary Figure 2). Thus, these bacteria were likely responsible for mediating the sulfate reduction detected.

There is little known about the anaerobic pathways for hydrocarbon degradation, especially in archaea. Comparison of MAGs to two databases of known hydrocarbon degradation proteins^22,23^. Most of the MAGs (81/93) had the metabolic potential to degrade hydrocarbons. The two novel ANME-2c in particular may be able to degrade long chain alkanes and naphthalene independently, as they encode several essential proteins involved in both aerobic and anaerobic hydrocarbon degradation (Supplementary Tables 7 and 8). Both contain genes for alkyl hydroperoxide reductase (ahpF) and an alkane hydroxylase system marker protein (alkH_ald) from the aerobic alkane degradation database^22,23^ as well as benzene carboxylase (AbcA_1 and AbcA_2), alkene C2 methylene hydroxylase (AhyA), flavin binding monooxygenase (AlmA), naphthalene carboxylase (K27540), long-chain alkane hydroxylases (LadA_alpha, LadB), Toluene-4-monooxygenase (TmoA_BmoA), and Toluene-ortho-monooxygenase (TomA3) proteins from the Calgary approach^22,23^. Additionally, CH_155 encodes two copies alkane hydroxylase system (alkN_mcp) for hydrocarbon sensing and CH_159 encodes three of the five alkane hydroxylase system biterminal oxidation rubredoxins (alkG_rub_rdx). Several of these hydrocarbon degradation marker genes have, to our knowledge, never been verified in archaea before.

It is possible that the bacteria in the community, like Desulfobacterota, Bacteroidota, Caldatribacterota, or Chloroflexota, are degrading oil in these enrichments and then passing hydrogen and carbon dioxide or acetate to ANME for the subsequent production of methane from less complex organic carbon compounds. We identified several other members of the enrichments that are capable of hydrocarbon degradation. For example, 7 other community members across diverse taxa including Thermoplasmatota, Thermotoga, Caldatribacterota, Desulfobacterota, Bacteroidota, Acidobacteria, and Chloroflexota) encoded 3 or more of the 6 alkane monooxygenase subunits (AlkBGHJKT) which are involved in the aerobic oxidation of alkanes to alcohols^24^ (Supplementary Figure 3 and Supplementary Table 9). Ten other MAGs, besides the 2 ANME-2c, encode naphthalene carboxylase (K27540) for degradation, based on searches using the CANT-HYD^23^ database (Supplementary Table 7). Thus, it is possible that there are metabolic handoffs acetate, CO_2_, and H_2_ from other community members to convert hydrocarbons to methane (Figure 4).

**Figure 4.**
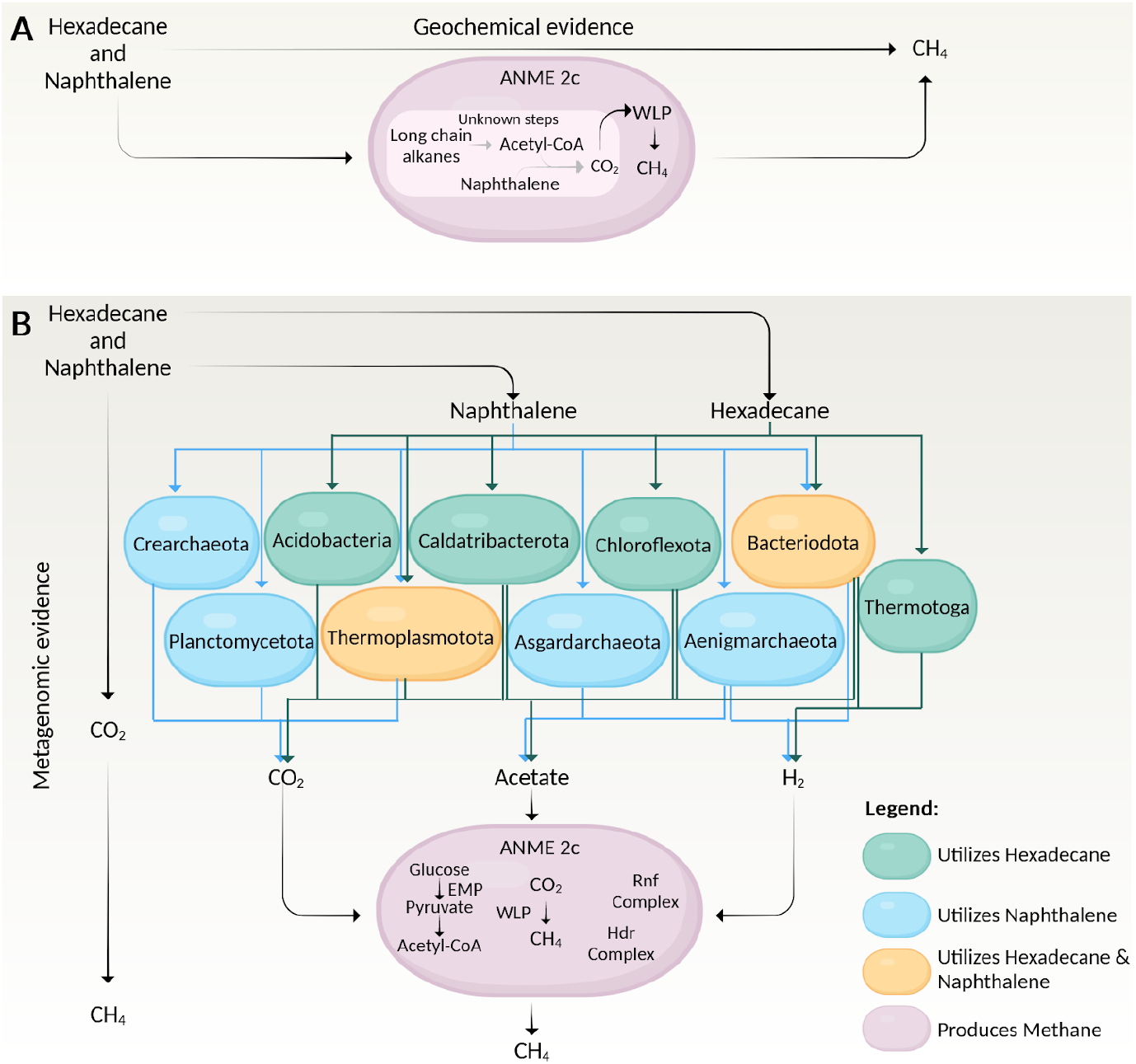
Carbon flow from hexadecane and naphthalene to methane in microbial communities of hydrothermally impacted sediments. Predicted flow of carbon from hexadecane and naphthalene to methane using geochemical (A) and metagenomic (B) evidence. Created with BioRender.com.

## Discussion

ANME archaea are broadly recognized to be mediating anaerobic oxidation of methane in marine sediments^25-26^. However, the radiotracer data suggest that radio labeled hydrocarbons (naphthalene and hexadecane) are degraded either syntrophically to CO_2_ and H_2_ or Acetate then passed to methanogens, or directly by reductive demethylation to produce methane. Traditional anaerobic oxidation of methane (AOM) pathways that ANME encodes, like most enzymatic reactions, can experience “back flux” when products build up resulting in “reversal” of the metabolic pathway without energy production^27^. This back flux is unlikely in these enrichment cultures as there was a media amendment (addition of crude oil) resulting in a high concentration of organic matter. Moreover, ANME-2c were only abundant microbes in our enrichment cultures that encode methane producing pathways. Therefore, ANME-2c appears to be producing methane directly from hydrocarbons or in very close syntrophy with another hydrocarbon degrader in the enrichments.

The cycling of methane by archaea on the ocean floor is responsible for consuming 90% of the methane fluxing through marine sediments, meaning that processes mediated by archaea are critical for understanding and predicting future climate. ANME archaea have not been definitively demonstrated to produce methane outside of the enzymatic backflux mechanism. Here we show that consortia, dominated by ANME-2c, from the deep sea are capable of degrading oil components to methane. Our findings suggest that like other low energy metabolic processes^28,29^, AOM by ANME-2c is reversible under certain conditions, perhaps in oil-rich environments. In order to predict the physiological activity of ANME in nature, a detailed examination of their energetic constraints is needed. These findings suggest that the biogeochemical roles of globally-distributed archaea may be more variable and dynamic than previously recognized. Their involvement in hydrocarbon degradation is unexpected and important, making them important sources of methane in the oceans, as well as important methane sinks.

## Methods

### Site description and sample collection

Sediment samples were collected during two research expeditions. The first to Guaymas Basin in the Gulf of California were collected with human-occupied vehicle *Alvin* during expedition AT42-05 onboard *R/V Atlantis* in November 2018. Two sites were sampled during AT42-05: Cathedral Hill (27°00.696’N, 111°24.365’W) and AcetoBalsamico (27°00.470’N, 111°24.427’W)^19,30^. The second expedition, FK190211, onboard *R/V Falkor* also sampled the Cathedral Hill site in Guaymas Basin using remotely operated vehicle *SUBastian* during February and March of 2019.

Sediment push cores were collected with HOV *Alvin* (AT42-05) or ROV *SUBastian* (FK190211). For all sites and expeditions, paired cores were sectioned on board immediately upon recovery for geochemical analysis, radiotracer incubations, and molecular analyses. During the expedition FK190211 paired push cores were collected using ROV *SuBastian* during dive 244 from the Cathedral Hill site and upon recovery on deck were stored under anoxic conditions at 4°C for subsequent enrichment experiments.

Sedimentary pore-fluids were extracted using a mechanical squeezer and filtered through a 0.2 μm syringe filter^31^. Subsamples for determination of porewater dissolved organic carbon (DOC) and nutrients were frozen and stored at −20 °C for shore-based analysis. Samples for sulfate analysis were preserved by addition of nitric acid (10 µL concentrated nitric acid per mL sample) and hydrogen sulfide (H_2_S) samples (unfiltered) were fixed with 20% zinc acetate. For each depth horizon, a 3 mL sediment sub-sample to determine dissolved methane concentration was transferred into a 20 mL serum vial containing 3 mL He-purged 2 M NaOH solution. Each vial was immediately crimp sealed, vortexed to halt biological activity, and stored at room temperature until analysis. For radiotracer incubations, ∼3 mL sediment was transferred into a modified cut-end Hungate tube as previously described^31,32^. Triplicate live samples from each sediment horizon and at least one killed control per core were taken for the quantification of rates for sulfate reduction, AOM, and hydrocarbon (hexadecane and naphthalene) turnover to methane. Sediments samples for metagenomic sequencing were frozen at -80°C for later nucleic acid extraction.

### Oil degrading enrichments

Back in the lab, the top layer of two cores (0-5 cm) from FK190211 dive 244 were homogenized in an anoxic glove box (N_2_/H_2_ 95/5%). The sediment was divided into two equal parts by mass and diluted in equal parts anoxic artificial medium to form a slurry^31–33^. One half of the slurry was made “sulfate free” (*i*.*e*., no sulfate was added to the medium), while the remaining slurry was amended to a final sulfate concentration of ∼28 mM. The resulting slurries were divided into 150 mL aliquots in 160 mL serum bottles (n=8 for each slurry type). Crude oil (225 µL) was added to each bottle to serve as a carbon source. For each treatment triplicate bottles were incubated at 4°C and 30°C to reflect approximate minimum and maximum temperatures detected from the *in situ* collections. The two remaining bottles for each treatment were autoclaved and served as kill controls. Each serum bottle was sealed with a blue butyl rubber stopper and aluminum crimp cap, and the headspace was purged for several minutes with 10% CO_2_ in N_2_ to remove excess H_2_ from the anaerobic chamber.

Microbial activity was assessed by measuring dissolved sulfide concentrations in each incubation vial. Once sulfide concentrations stopped increasing (highest observed concentration was ∼12 mM) subsamples were collected to quantify sulfate reduction and methane production rates and nucleic acids were extracted.

### Geochemical analysis

Dissolved methane concentrations were measured via a gas chromatograph (SRI) equipped with a flame ionization detector (GC-FID) using ultra high purity methane standards (Airgas). Porewater sulfate was quantified using a Dionex^®^ Ion Chromatograph. A standard curve was generated from the IAPSO seawater standard (P series). DOC concentrations were determined with a Shimadzu TOC-V equipped with a nondispersive infrared detector. Reagent grade potassium hydrogen phthalate was used as a standard for the analysis. Porewater hydrogen sulfide concentrations in field samples and in the enrichments were measured spectrophotometrically via the Cline assay^34,35^. During the expedition FK190211 *in situ* temperature profiles were collected manually with the heat flow probe on the ROV *SuBastian*.

### DNA extraction and Metagenome assembly

Samples for DNA analyses were collected in sterile cryotubes (Nalgene), flash frozen in liquid nitrogen, and stored at -80ºC. Sedimentary DNA then extracted using a Qiagen DNeasy PowerMax Soil kit with additional bead beating steps. DNA was quantified using a Qubit and sent to CosmosID (Germantown, MD)^36^ for sequencing. Metagenomic sequencing library preparations followed the manufacturer’s protocol (VAHTS Universal DNA Library Prep Kit for Illumina). For each sample, 100 ng of genomic DNA was randomly fragmented to <500 bp by sonication (Covaris S220). The fragments were treated with End Prep Enzyme Mix for end repairing, 5’ phosphorylation and dA-tailing in one reaction, followed by a T-A ligation to add adapters to both ends. Size selection for Adapter-ligated DNA was then performed using VAHTSTM DNA Clean Beads, and fragments of ∼470 bp (with the approximate insert size of 350 bp) were recovered. Each sample was then amplified by PCR for 8 cycles using P5 and P7 primers, with both primers carrying sequences which can anneal with the flow cell to perform bridge PCR and P7 primer carrying a six-base index which enabled multiplexing. The PCR products were cleaned using VAHTSTM DNA Clean Beads, validated using an Agilent 2100 Bioanalyzer (Agilent Technologies, Palo Alto, CA, USA), and quantified by Qubit 3.0 Fluorometer (Invitrogen, Carlsbad, CA, USA). Then libraries with different indices were multiplexed and loaded on an Illumina HiSeq instrument according to manufacturer’s instructions (Illumina, San Diego, CA, USA).

We then assembled the 64 Gb of read-read shotgun metagenomic sequencing data from these enrichments. Reads were trimmed for quality and adapters using Trimmomatic v0.39^37^ with the following parameters: leading:5, trailing:5, and sliding window: 5:15. Reads that were shorter than 50 base pairs were removed. TruSeq adapters were then removed as follows TruSeq2-PE.fa:2:30:10:8:True. Interleaving of high quality paired end reads was completed with BBMap v39.01^37,38^. Interleaved reads were then assembled with IDBA v1.1.3^39^ with the following parameters; pre correction, mink 65, maxk 115, step 10, seed_kmer 55. Coverage was obtained by mapping all interleaved reads of each sample against the assembly using the BWA-MEM algorithm v0.1.5.1^40^ in paired-end mode. Only contiguous sequence segments (contigs) larger than 2,000 base pairs were utilized in the next stages of assembly.

Contigs from the IDBA assembly were co-binned using MetaBAT v2.12.1^40,41^ using the following parameters: --minCVSum 0 --saveCls -d -v --minCV 0.1 -m 2000 and CONCOCT v1.1.0^42^ using the following parameters: --clusters 400 --kmer_length 4 --length_threshold 3000 --seed 4 --iterations 500. The resulting metagenome assembled genomes (MAGs) were combined from these two binning workflows using DAS Tools v1.1.2^42,43^. For both MetaBAT and CONCOCT, a scaffold-to-bin list was prepared, and DAS Tools was run on each of the eight scaffold files with these parameters: DAS_Tools -i Concoct.scaffolds.tsv, Metabat.scaffolds.tsv -l concoct,metabat -c assembly.contigs.fasta –debug -t –write_bins 1 –search_engine_blast. The accuracy of all MAGs was evaluated by calculating the percent completeness and gene duplication using the CheckM lineage workflow v1.0.5^42–44^. Only MAGs that were both 50% or more complete and contained 10% or less gene duplications were included for further analyses. Relative abundance of each MAG was calculated using CoverM v0.6.1^11^.

### Taxonomy and phylogenetic reconstruction

GTDB-Tk classify workflow v0.3.3^10^ was used for taxonomic identification of each MAG. This workflow consisted of three steps; identify, align, and classify. During the GTDB-Tk identify step the MAGs were divided into 77 bacteria and 16 archaea using Prodigal^45,46^ and HMMER^47^ to identify 120 bacterial and 53 archaeal marker genes^45^. Then for each MAG the align and classify steps were implemented independently first on the bacterial dataset and then the archaeal dataset. The align step concatenated the aligned marker genes and filtered based on a 5,000 amino acid cutoff. Finally, the classification step employed pplacer^48^ which uses maximum-likelihood to place each MAG in the GTDB-Tk reference tree^49^. GTDB-Tk then assigned taxonomy by combining placement in the GTDB-TK reference tree and relative evolutionary divergence (RED) values with reference genomes.

An archaeal phylogeny was generated to verify the GTDB-Tk classification of these enrichment genomes using 37 conserved marker genes that are mainly ribosomal. These 37 markers were extracted from the 16 archaeal MAGs using Phylosift v1.0.1^50^. We used one of the 37 marker proteins identified to perform a BLASTP^16^ search against the NCBI RefSeq database^51^ to obtain the publicly available genomes that are most closely related to the ANME 2c metagenomes of interest. For these reference genomes, we then extracted the 37 marker genes using Phylosift and added them to the analyses. Alignments of the extracted assembled MAGs and reference genomes were generated using MAFFT v7.310^52^ with the following parameters: –maxiterate 1000 –localpair. A list of the publicly available reference genomes used for both the archaeal and bacterial phylogenies can be found in the supplementary data (Supplementary Table 10). The MAFFT alignment was then trimmed using ClipKit^53^ with the following parameters: -m gappy. Then the phylogeny was constructed with IQTree v1.6.12^54^ as follows: -bb 1000 -bnni and the LG+F+R10 model. This phylogenetic reconstruction was also performed for the 77 bacterial MAGs using Phylosift v1.0.1^50^, MAFFT v7.3.10^52^, and IQTree v1.6.12^54^ as detailed above except with the LG+R10 model. Both the archaeal and bacterial phylogenies were visualized using iTOL v6^55^.

### Phylogenetic reconstruction of McrA

1,216 McrA protein sequences^21^ were used to generate a DIAMOND database to conduct DIAMOND BlastP v2.1.8.162^20^ searches against the 93 enrichment genomes to determine which community members were capable of methane production. Only BlastP^16^ hits with greater than 60% identity and over 200 base pairs in length were considered in downstream analyses. This BlastP^16^ search resulted in isolation of two McrA sequences from the assembled enrichment genomes, one from bin CH_155 and the other from bin CH_159 which were both identified as ANME group 2c archaea. These two McrA sequences were then aligned with the 1,216 reference sequences using MAFFT v7.310^56^ with the following parameters: –maxiterate 1000 –localpair. The MAFFT alignment was manually trimmed and gaps over 10% were masked in Geneious Prime 2023.2.1. Then the phylogeny was constructed with IQTree v1.6.12^54^ using the LG+F+R6 model as follows: -bb 1000 -bnni. The phylogeny was visualized using iTOL v.6^55^.

### Phylogenetic reconstruction of alpha (DsrA) and beta (DsrB) dissimilatory sulfate reductase subunits

Manually curated dissimilatory sulfate reductase (DsrA and DsrB) protein reference databases of 1,505 and 1,695 sequences respectively were used to generate two independent DIAMOND databases to perform DIAMOND BlastP v2.1.8.162^20^ searches against the 93 enrichment genomes to determine which community members were capable of sulfate reduction. This BlastP^16^ search resulted in isolation of 12 DsrA and 13 DsrB sequences from the assembled enrichment genomes (Supplementary Figure 2). These sequences were then aligned (independently for DsrA and DsrB) with the corresponding reference sequences using MAFFT v7.310^56^ with the following parameters: –maxiterate 1000 –localpair. The two MAFFT alignments were manually trimmed and gaps over 10% were masked in Geneious Prime 2023.2.1. Then the phylogenies were independently constructed with IQTree v1.6.12^54^ using the LG+R10 model for DsrA and the LG+R9 model for DsrB as follows: -bb 1000 -bnni. The phylogenies were visualized using iTOL v.6^55^.

### Phylogenetic reconstruction of nickle-iron (NiFe) and iron-iron (FeFe) hydrogenases

Two Diamond databases from 2,015 NiFe and 1,221 FeFe reference sequences were created to perform DIAMOND BlastP v2.1.8.162^20^ searches across all 93 metagenomes. This BlastP^20^ search resulted in identification of 2,310 potential NiFe sequences and 184 potential FeFe sequences. These sequences were then filtered by bitscore according to the cutoffs established in HydDB^57^ and by CxxC motif presence to yield 106 NiFe sequences across 58 metagenomes (Supplementary Figure 4) and 69 FeFe sequences across 33 metagenomes (Supplementary Figure 5). These sequences were then aligned (independently for NiFe and FeFe hydrogenases) with corresponding reference sequences using MAFFT v7.310^56^ with the following parameters: –maxiterate 1000 –localpair. The two MAFFT alignments were manually trimmed and gaps over 10% were masked using ClipKit v.x.x.x. Then the phylogenies were independently constructed with IQTree v1.6.12^54^ using the LG+R10 model for NiFe and the LG+R10 model for FeFe as follows: -bb 1000 -bnni -safe. The phylogenies were visualized using iTOL v.6^55^.

### Community metabolic reconstruction

Predicted proteins from Prodigal translation of nucleic acid sequences for individual MAGs were characterized using several well-known databases: KofamScan^58^, MEBS^59^, CANT-HYD^23^, HADEG^60^, and 38 conserved methanogenesis marker genes^21^. Searches over these databases were performed using default parameters. A subset of data from supplementary table 5 (taking the top 5 most abundant bacteria and archaea) served as the input file to generate the bubble plot in figure 3 above with the count of marker genes for each genome across the methanogenesis marker genes databases aforementioned^21^ (Supplementary Table 5). CANT-HYD^23^ and HADEG^60^ annotations (Supplementary Tables 7 and 8) were used to predict which microbial community members were likely able to degrade hydrocarbons (hexadecane and naphthalene) while MEBS^59^ (Supplementary Table 12) calculations were used to predict general metabolic inputs and outputs for all 93 community members (Fig. 4).

Alkane monooxygenases were identified in all 93 MAGs from manually curated databases of 395 AlkB, 52 AlkG, 796 AlkH, 236 AlkJ, 763 AlkK, and 76 AlkT reference sequences. Proposed AlkB DIAMOND BlastP v2.1.8.162^20^ hits were compared to both KEGG^58^ and BLAST-P^16^ annotations for all scaffolds. Scaffolds that were annotated with KEGG and matched BLAST-P hits for similar protein families that were annotated with the manually curated alkane monooxygenase (AlkB) database, were then aligned to the 395 reference sequences to generate a phylogeny as described for the McrA tree (Supplementary Figure 3). 14 MAGs were found to contain three of the six (AlkBGHJKT) alkane monooxygenase subunits.

CANT-HYD searches^23^ were conducted below the noise cutoff because none of the 93 MAGs had any hits to this database with the trusted or noise default cutoffs. Then the proposed hits from the CANT-HYD and HADEG^60^ databases were used to predict community members capable of degrading hexadecane and/or naphthalene. Together these six databases: KEGG^58^, MEBS^59^, 38 methanogenesis marker genes^21^, alkane monooxygenase references^24^, CANT-HYD^23^, and HADEG^60^ were used to generate the predicted enrichment community hydrocarbon degradation and methane production capabilities (Figure 4).

### ANME Group 2c (*Ca. Methanogaster*) metabolic reconstruction

Predicted genes for the two ANME-2c MAGs were also characterized using these same aforementioned databases: KofamScan^58^, MEBS^59^, HADEG^60^, CANT-HYD^23^ and 38 conserved methanogenesis marker genes^21^. KEGG^58^ and MEBS^59^ annotations (Supplementary Tables 12 and 13) were used to reconstruct the ANME-2c metabolic cell diagram in figure 4. A more detailed metabolic diagram for the ANME-2c MAGs can be found in the supplementary information (Fig. S6) KEGG searches conducted with KofamScan^58^ demonstrated that these ANME-2c MAGs have three missing steps in methanogenesis as aforementioned. However, these three steps; *fpo, frh*, and *hdrDE* have previously been shown to be missing from several ANME-2c genomes before^18^ and would be involved in the traditional anaerobic oxidation of methane metabolic processes in ANME.

## Acknowledgments

This research was supported by an Investigator award from the Simons Foundation Award LI-SIAME-00002001 to B.J.B. and by the National Science Foundation’s Biological Oceanography program through awards OCE-1357360 and OCE-2049439 (to S.B.J.). We thank the captain and crew of R/V Atlantis and the HOV Alvin team and the shipboard science party for their help securing and processing samples during expedition AT42-05. We thank the Schmidt Ocean Institute for providing time at sea on R/V Falkor. We thank the captain and crew of R/V Falkor, the ROV SuBastian team, and the shipboard science party for their assistance with sample collection and processing during expedition FK190211. We also thank Dr. Mirna Vasquez Rosas Landa, now at the Universidad Nacional Autónoma de México, for genomic assembly and binning as well as Zachary Marinelli and Kimberly S. Hunter from the University of Georgia, Athens for their assistance with geochemistry measurements.

## Author Contributions

N.S. reconstructed MAGs. N.S., B.R.M. and V.D.A. performed metabolic reconstructions. A.M., G.C.Z., S.B.J. and B.J.B. collected the samples. A.M., G.C.Z and Z.M. performed enrichment experiments. Geochemical measurements were performed by A.M. N.S. and B.J.B. wrote the manuscript, and all authors provided comments on the manuscript. S.B.J. and B.J.B. conceived and advised the research.

## Data availability

All metagenome sequences are available from the National Center for Biotechnology Information (NCBI) Read Archive under the biosample accession ID: PRJNA924812 and the submission metadata are in the supplementary information (supplementary table 1).

