## Supplementary Figures 1-6 for "Methane production from hydrocarbons by consortia dominated by ANME archaea"

#Corresponding authors

### Supplementary Figures:

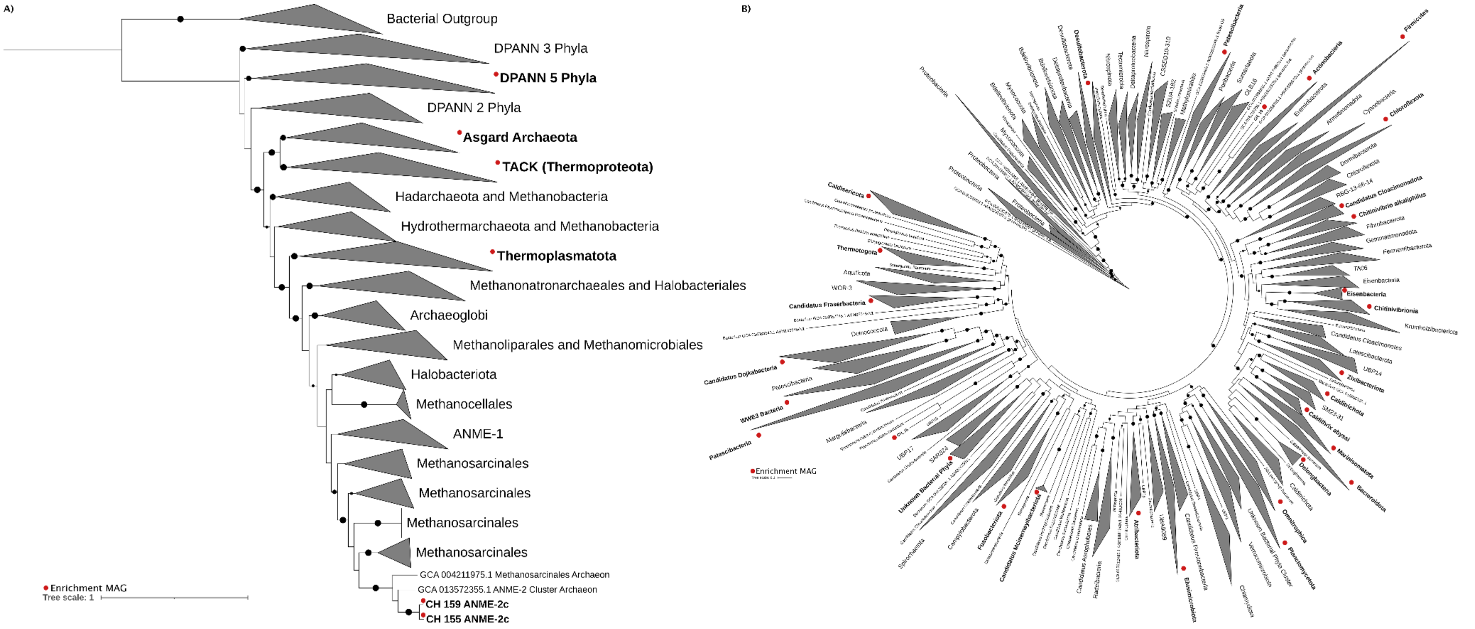


**Supplementary Figure 1. Phylogenetic position of enrichment MAGs reconstructed from the communities.** Black dots indicate a bootstrap value ranging from smallest to largest dots 90 to 100. Red dots indicate a lineage containing one or more MAG(s) from the enrichment community. (A) The archaeal phylogeny was generated using ribosomal proteins of 37 concatenated marker proteins[1] and 740 reference sequences (supp table with IDs). There were 16 archaeal MAGs across 9 phyla. (B) The bacterial phylogeny was generated using ribosomal proteins of 37 concatenated marker proteins[1] and 1,439 reference sequences (supp table with IDs). There were 77 bacterial MAGs across 25 phyla.

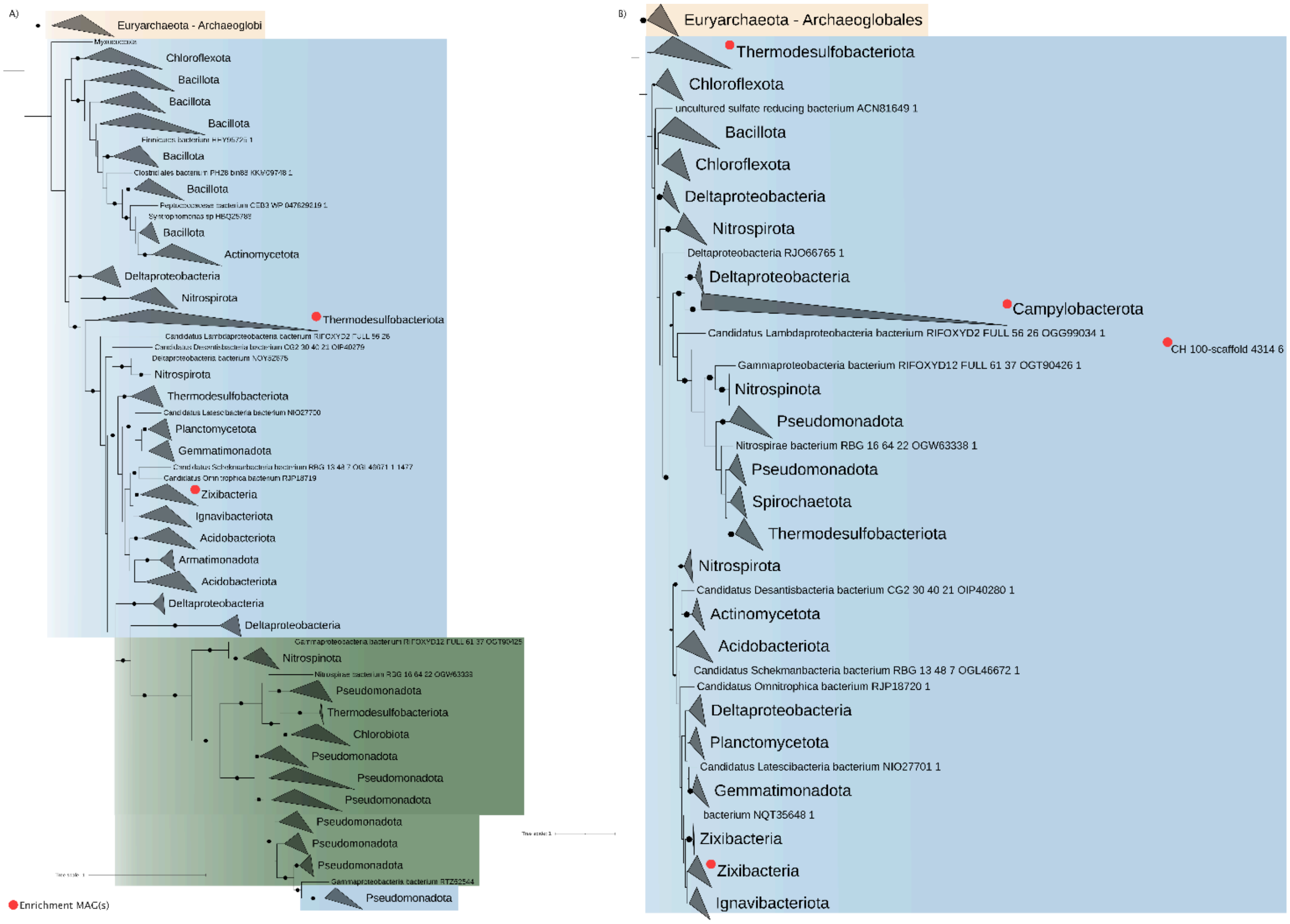


**Supplementary Figure 2. DsrA and DsrB phylogenetic trees.** Both trees were generated by searching across all 93 MAGs for 1,505 DsrA and 1,695 DsrB proteins with Diamond BlastP[2]. Red dots indicate one or more enrichment genomes with the DsrA or DsrB respectively. Black dots indicate a bootstrap value ranging from smallest to largest dots 90 to 100. Lineages highlighted in yellow are the outgroup Archaeoglobales, in blue are reductive Dsr proteins, and in green are oxidative Dsr proteins. (A) This phylogeny is the alpha subunit of the dissimilatory sulfate reductase (DsrA). Diamond BlastP[2] searches identified 11 DsrA in the enrichment community. 10 from Thermodesulfobacteriota and 1 from Zixibacteria. (B) This phylogeny is the beta subunit of the dissimilatory sulfate reductase (DsrB). Diamond BlastP[2] searches identified 13 DsrB in the enrichment community. 10 from Thermodesulfobacteriota, 1 from Campylobacterota, 1 from Firmicutes (CH_100), and 1 from Zixibacteria.

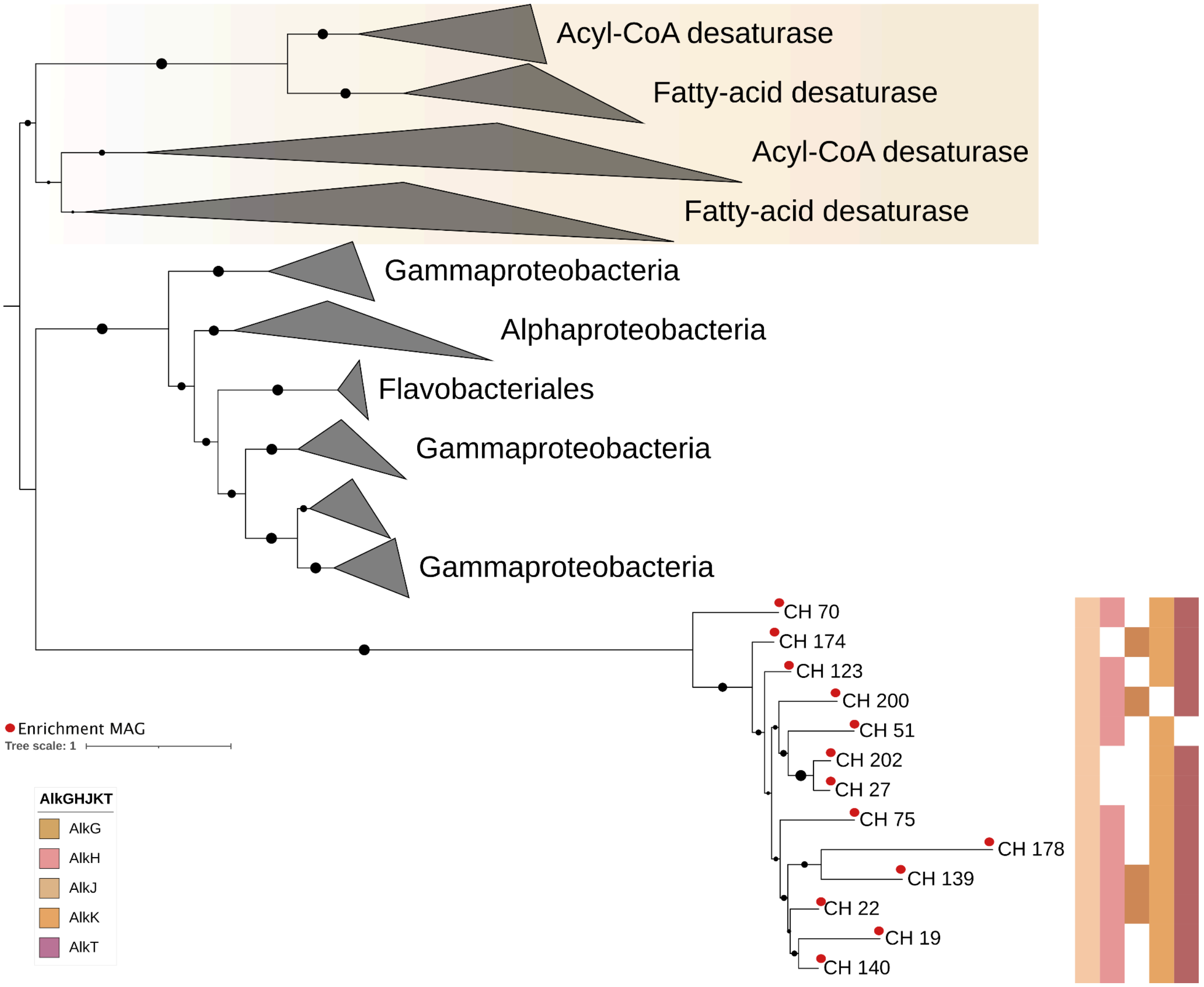


**Supplementary Figure 3. Phylogenetic position of predicted alkane monooxygenase (AlkB) proteins.** 13 MAGs encoded three or more subunits from the six total subunits (AlkBGHJKT). Red dots indicate enrichment MAGs. Colored strips to the right of the enrichment MAGs indicate additional alkane monooxygenase subunits encoded in each MAG from left to right AlkG, AlkH, AlkJ, AlkK, and AlkT. The tan highlighted region of the phylogeny is the outgroup. Black dots indicate a bootstrap value ranging from smallest to largest dots 90 to 100. For taxonomic information of enrichment MAGs see supplementary table 2.

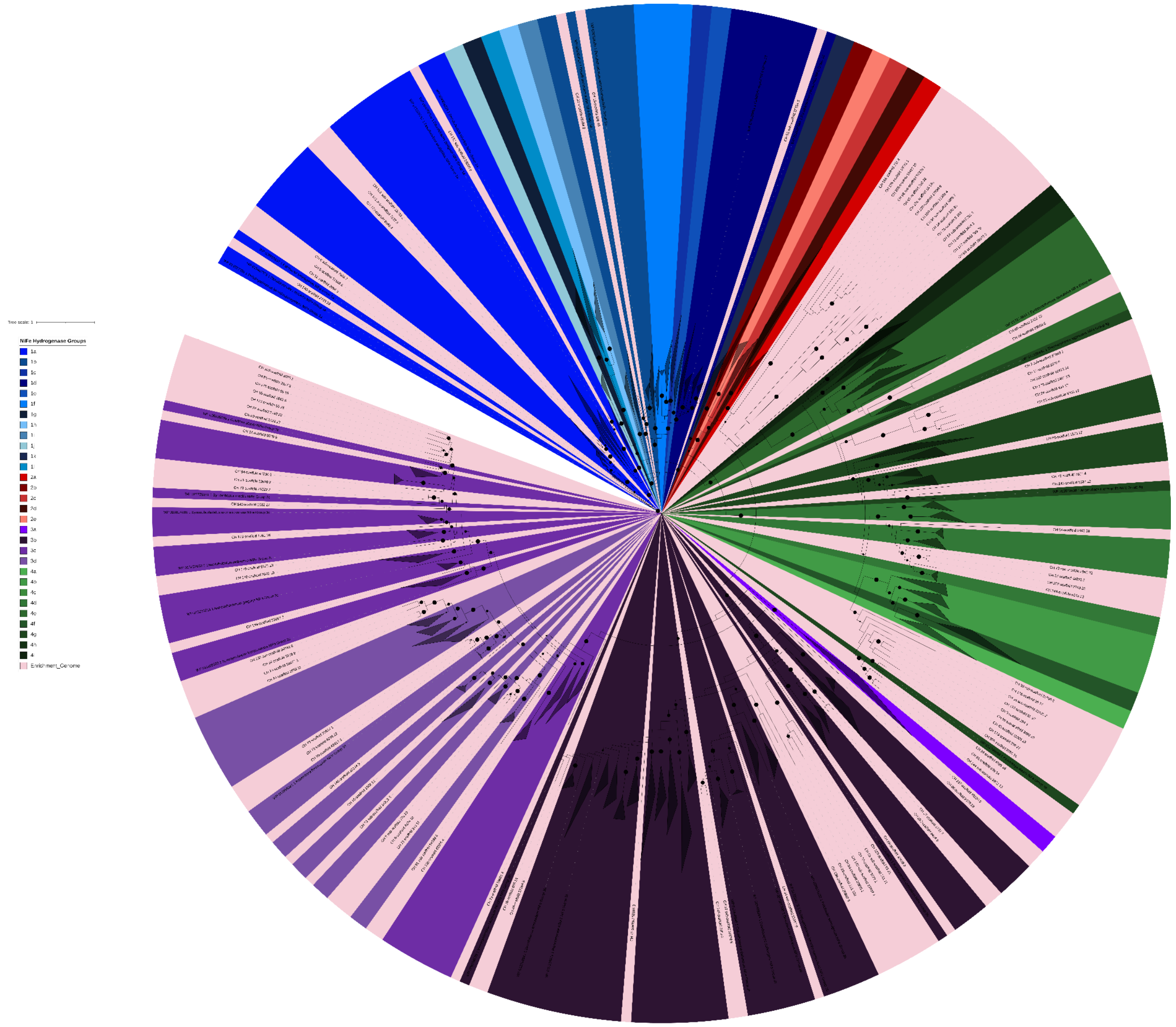

**Supplementary Figure 4. Nickel-iron hydrogenase phylogenetic tree.** 38 nickel-iron hydrogenase classes were used from HydDB[3] to conduct HMM searches[4] across all 93 enrichment MAGs. By applying bitscore and motif filtering[5] 106 nickel-iron hydrogenases were identified across 57 MAGs and are highlighted in pink in the tree.

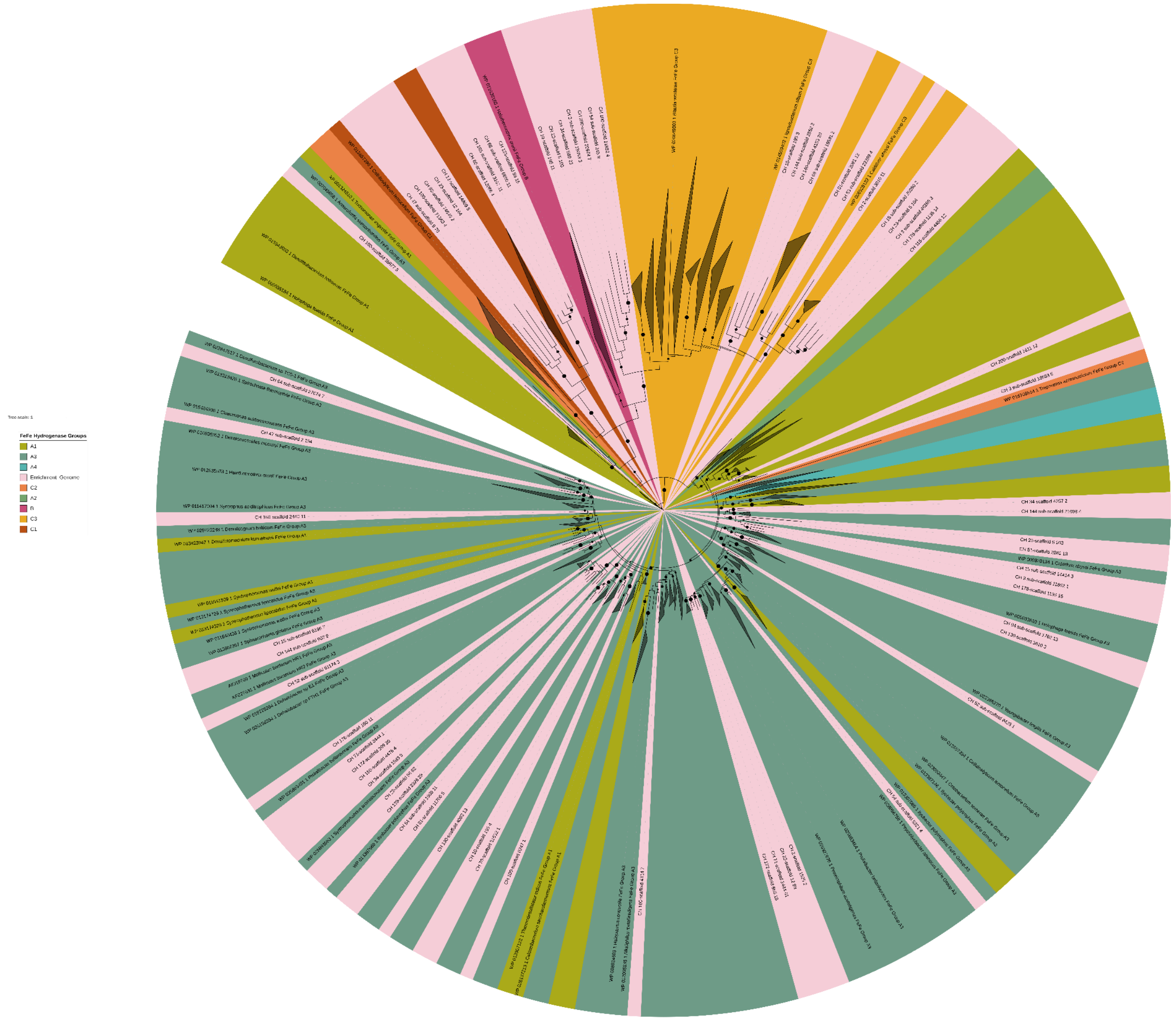


**Supplementary Figure 5. Iron-iron hydrogenase phylogenetic tree.** 8 iron-iron hydrogenase classes were used from HydDB[3] to conduct HMM searches[4] across all 93 enrichment MAGs. By applying bitscore and motif filtering[5] 69 iron-iron hydrogenases were identified across 33 MAGs which are highlighted in pink.

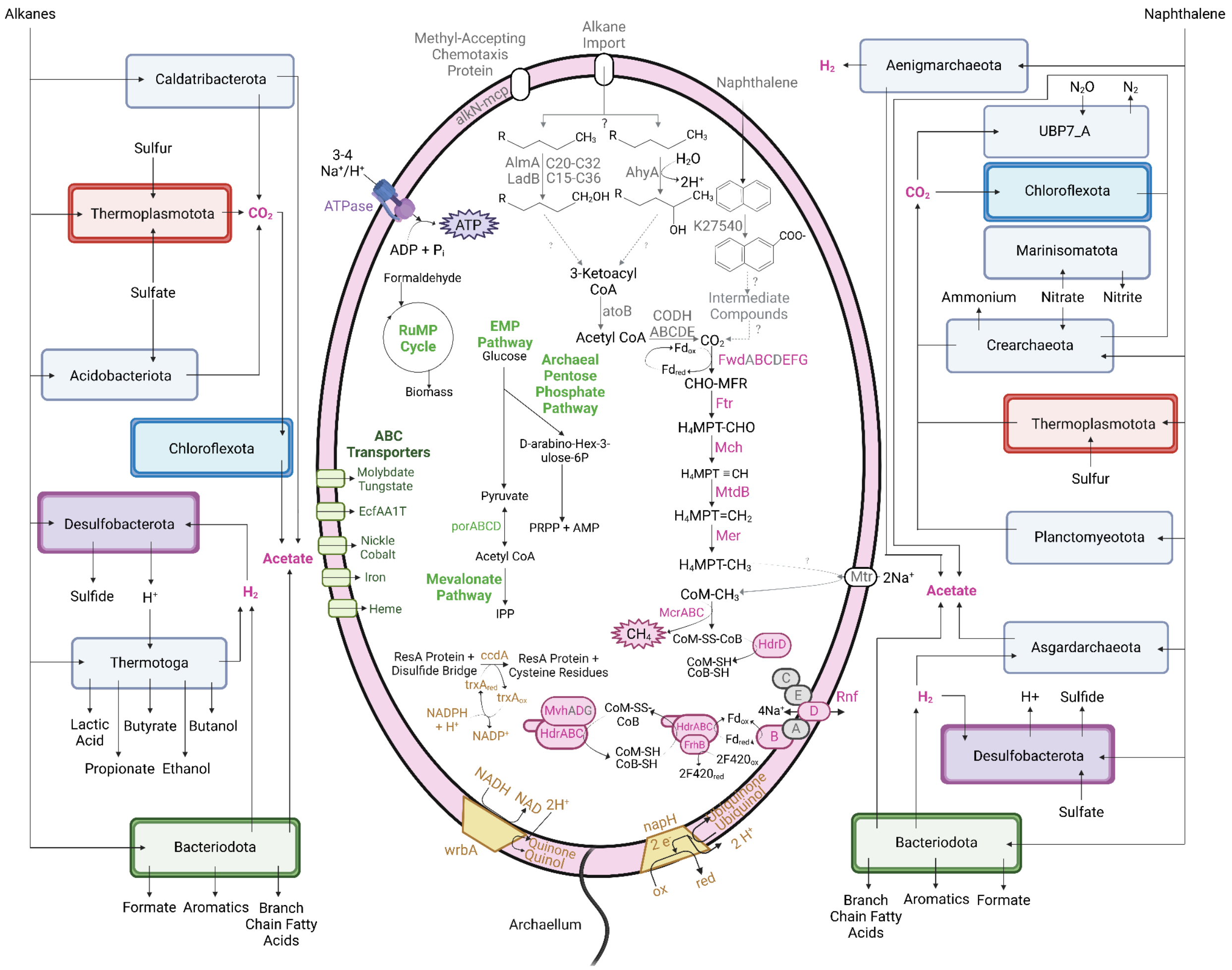


**Supplementary Figure 6. Predicted hydrocarbon degradation of the enrichment community and methanogenesis pathways in ANME-2c archaea.** The large cell in the center is the metabolic capabilities of the two ANME-2c enrichment genomes. The organisms to the left of the ANME cell are community members capable of alkane consumption while the organisms to the right may consume naphthalene. All of the black, solid arrows represent the presence of genes or proteins in the organisms. Dashed lines and everything in gray are missing steps of a potential metabolic pathway. Created with BioRender.com.
